# Linked supergenes underlie split sex ratio and social organization in an ant

**DOI:** 10.1101/2021.01.22.427864

**Authors:** German Lagunas-Robles, Jessica Purcell, Alan Brelsford

**Affiliations:** Department of Evolution, Ecology and Organismal Biology, University of California, Riverside, Riverside, CA 92521, USA; Department of Entomology, University of California, Riverside, Riverside, CA 92521, USA

## Abstract

Sexually reproducing organisms usually invest equally in male and female offspring. Deviations from this pattern have led researchers to new discoveries in the study of parent-offspring conflict, genomic conflict, and cooperation. Some social insect species exhibit the unusual population-level pattern of split sex ratio, wherein some colonies specialize in the production of future queens and others specialize in the production of males. Theoretical work focused on the relatedness asymmetries emerging from haplodiploid inheritance, whereby queens are equally related to daughters and sons, but their daughter workers are more closely related to sisters than to brothers, led to a series of testable predictions and spawned many empirical studies of this phenomenon. However, not all empirical systems follow predicted patterns, so questions remain about how split sex ratio emerges. Here, we sequence the genomes of 138 *Formica glacialis* workers from 34 male-producing and 34 gyne-producing colonies to determine whether split sex ratio is under genetic control. We identify a supergene spanning 5.5 Mbp that is closely associated with sex allocation in this system. Strikingly, this supergene is adjacent to another supergene spanning 5 Mbp that is associated with variation in colony queen number. We identify a similar pattern in a second related species, *Formica podzolica.* The discovery that split sex ratio is determined, at least in part, by a supergene in two species opens a new line of research on the evolutionary drivers of split sex ratio.

**Significance Statement:** Some social insects exhibit split sex ratio, wherein some colonies produce future queens and others produce males. This phenomenon spawned many influential theoretical studies and empirical tests, both of which have advanced our understanding of parent-offspring conflicts and cooperation. However, some empirical systems did not follow theoretical predictions, indicating that researchers lack a comprehensive understanding of the drivers of split sex ratio. Here, we show that split sex ratio is associated with a large genomic region in two ant species. The discovery of a genetic basis for sex allocation in ants provides a novel explanation for this phenomenon, particularly in systems where empirical observations deviate from theoretical predictions.

## Introduction

The relative investment in male versus female offspring is a vital fitness component of sexually reproducing organisms. Research on sex allocation theory has yielded breakthroughs in our understanding of topics as diverse as parent-offspring conflict, evolution of cooperation, and genomic conflict (1). Parent-offspring conflict is predicted to occur in subdivided populations with strong local mate competition, as seen in polyembryonic parasitoids (2), and in systems with relatedness asymmetry between sisters and brothers, as found in haplodiploid species such as the primitively eusocial wasp *Polistes chinensis antennalis* (3). Parental control of sex ratio is also thought to contribute to the maintenance of cooperative breeding; for example, Seychelles warblers living in high quality territories where helpers provide strong benefits produce an excess of females, the helping sex (4). However, similar patterns of biased sex allocation increasing the frequency of the helping sex are not found among all cooperatively breeding birds (5). The discovery of a chromosome that skews sex ratio from female-biased to 100% male in the jewel wasp *Nasonia vitripennis* was the first clear empirical illustration of an intragenomic conflict (6). This paternal sex ratio chromosome is transmitted through sperm to fertilized eggs, where it causes the loss of other paternally inherited chromosomes to produce exclusively male offspring (7, 8). Subsequent discoveries of sex ratio distorter systems take different forms, including female biased sex ratios mediated by endosymbionts (9, 10). These studies opened the door for additional research on intragenomic conflict in multiple contexts, including between sexes (11, 12) and between social insect castes (13).

Where there is intragenomic conflict, one resolution is evolution of suppressed recombination to reduce the frequency of deleterious multilocus genotypes. This is illustrated in the standard model of sex chromosome evolution (14, 15), in which selection favors the loss of recombination between a sexually antagonistic locus and a sex-determining locus on the same chromosome, eventually leading to a Y or W chromosome that is exclusively present in one sex. Under the “reduction principle” (16), this is also expected to occur around sex-ratio distorters. In line with this prediction, sex-ratio distorter loci often occur in regions of low recombination (17–20), but we lack evidence for the direction of causality. The reduction principle is also expected to contribute to the formation of autosomal supergenes controlling other complex traits that involve epistatic interactions between two or more loci. Such supergenes have been found to control phenotypes including polymorphic wing coloration in butterflies, mating strategies in birds and fungi, self-incompatibility in plants, and colony social organization in ants (21–28). Autosomal supergenes, like sex chromosomes, are likely to represent the resolution of past intragenomic conflict between two or more loci.

Supergenes underlie at least two independently evolved cases of social polymorphism in ants. In the fire ant *Solenopsis invicta,* colony queen number is controlled by a supergene spanning most of a single chromosome (27). *Formica selysi* has a similar chromosome-spanning supergene underlying colony queen number, but there is no detectable overlap in gene content between the two (28). More recently, both ant social supergenes were shown to underlie colony queen number in other congeneric species (29, 30). In both systems, the haplotype associated with multi-queen (= polygyne) social structure is a selfish transmission distorter (31–33). These discoveries raise new questions about links between social structure and sex ratio that have been proposed in classic literature about sex allocation in Hymenoptera.

Trivers and Hare (34) proposed that queen-worker conflict, which is shaped by relatedness asymmetry within each nest, drives biased sex ratios. Since workers are more related to their full sisters (average relatedness = 0.75) than to their brothers (average relatedness = 0.25), workers in single-queen, monandrous colonies should favor the production of queens over males. Trivers and Hare (34) predicted that worker interests would prevail in these cases, resulting in female-biased offspring production. Queens are equally related to male and female offspring, so they should generally favor a 1:1 sex ratio. In colonies with multiple queens or multiple mates, the low relatedness reduces this conflict between queens and workers, resulting in weaker selection for biased sex allocation (34). Although these predictions revolutionized the way that researchers think about fitness and relatedness in social insect colonies, they are not ubiquitously upheld in empirical studies (1).

Strikingly, some social insect species exhibit a nearly complete segregation of male and queen production at the colony level, in a phenomenon known as ‘split sex ratio’. This extreme case has been observed in at least 20 different genera of ants, wasps, and bees (35–37). Boomsma and Grafen (36) argued that this pattern is consistent with worker control of sex ratio in populations with variation in relatedness asymmetry: workers that are more related than the population average to their nestmates should favor specializing in the production of new queens (hereafter “gynes”), while those that are less related than average should specialize in male production (36, 38). The variation in relatedness asymmetry would emerge from the number of mates per queen and from the number of queens per colony.

The models of Boomsma and Grafen inspired a burst of empirical research on split sex ratio in ants and other social insects (37, 39–45, 45–53). Ants in the genus *Formica* emerged as a prominent model system, as a result of their widespread and well documented variation in sex ratio and social structure (35). Many species exhibit split sex ratio or highly biased sex ratio (34, 41–45), but not all of these examples follow predicted patterns based on relatedness asymmetry. Finnish populations of *Formica truncorum* and *F. exsecta* follow theoretical predictions: in colonies with a single queen (= monogyne), monandrous queens tend to produce gynes, while polyandrous queens tend to produce males (41, 44, but see 46). A similar pattern was found in monogyne and polygyne colonies in *F. truncorum* (with polygyne colonies producing males (45)). A socially polymorphic population of *F. selysi* and a polygynous population of *F. exsecta* that exhibits variation in relatedness asymmetry deviated from these predicted patterns (46, 48). Additional studies have identified potential roles of habitat and diet in shaping sex allocation in *F. podzolica* (44), in *F. exsecta* (49), in *F. aquilonia* (50), as well as colony needs for queen replacement (51, 55). Finally, although *Wolbachia* is present in some *Formica* species exhibiting split sex ratio, it does not appear to influence sex ratio in any system studied so far (52, 53).

Taken together, it appears that there are yet missing pieces to the puzzle of how and why ants achieve a split sex ratio. A meta-analysis attributed only ~25% of the observed variance in sex allocation to relatedness asymmetry and variation in queen number (37). Theoretical examinations following from this finding support a possible role for virgin queens (which would produce only male offspring) or queen replacement (56), but another possibility is that *sex allocation by queens is itself under genetic control.*

Here, we examine the evidence for this mechanism, which could be responsible for much of the unexplained variance in patterns of split sex ratio. We 1) conduct a genome-wide association study for variants associated with sex ratio in *Formica glacialis,* 2) infer transmission patterns of sex-ratio-associated variants from colony-level genotype frequencies, 3) evaluate whether sex ratio and social organization map to the same region of the genome, and 4) test for a shared genetic basis of sex ratio in the related species *F. podzolica*.

## Results

Through a genome-wide association study (GWAS) of 138 *F. glacialis* whole-genome sequences, we identify numerous variants associated with colony sex allocation in a region of chromosome 3 spanning 5.5 Mbp (Figure 1a). A principal component analysis (PCA) of variants on chromosome 3 reveals three distinct genotype clusters, one of which is observed in just six individuals (Fig 1b). Of the workers with low PC2 scores (yellow and green clusters, Fig. 1b) 60.2 % were collected from male-producing colonies, while 93.3% of workers with high PC2 scores (purple cluster, Fig. 1 b) were from gyne-producing colonies. We investigate the parts of chromosome 3 that distinguish the genotype clusters through an assessment of F_ST_ between the clusters (Fig. 2). This comparison reveals two adjacent regions of differentiation. Between the two clusters with low PC1 scores, we observe differentiation spanning the region from 2-7.5 Mbp (Fig. 2a), similar to the region revealed in the initial GWAS. Between the two clusters with low PC2 values (both of which harbored an excess of workers from male producing colonies), we identify a differentiated region from about 7.5-12.5 Mbp, as well as a small peak at 2 Mbp (Fig. 2b).

**Figure 1.**
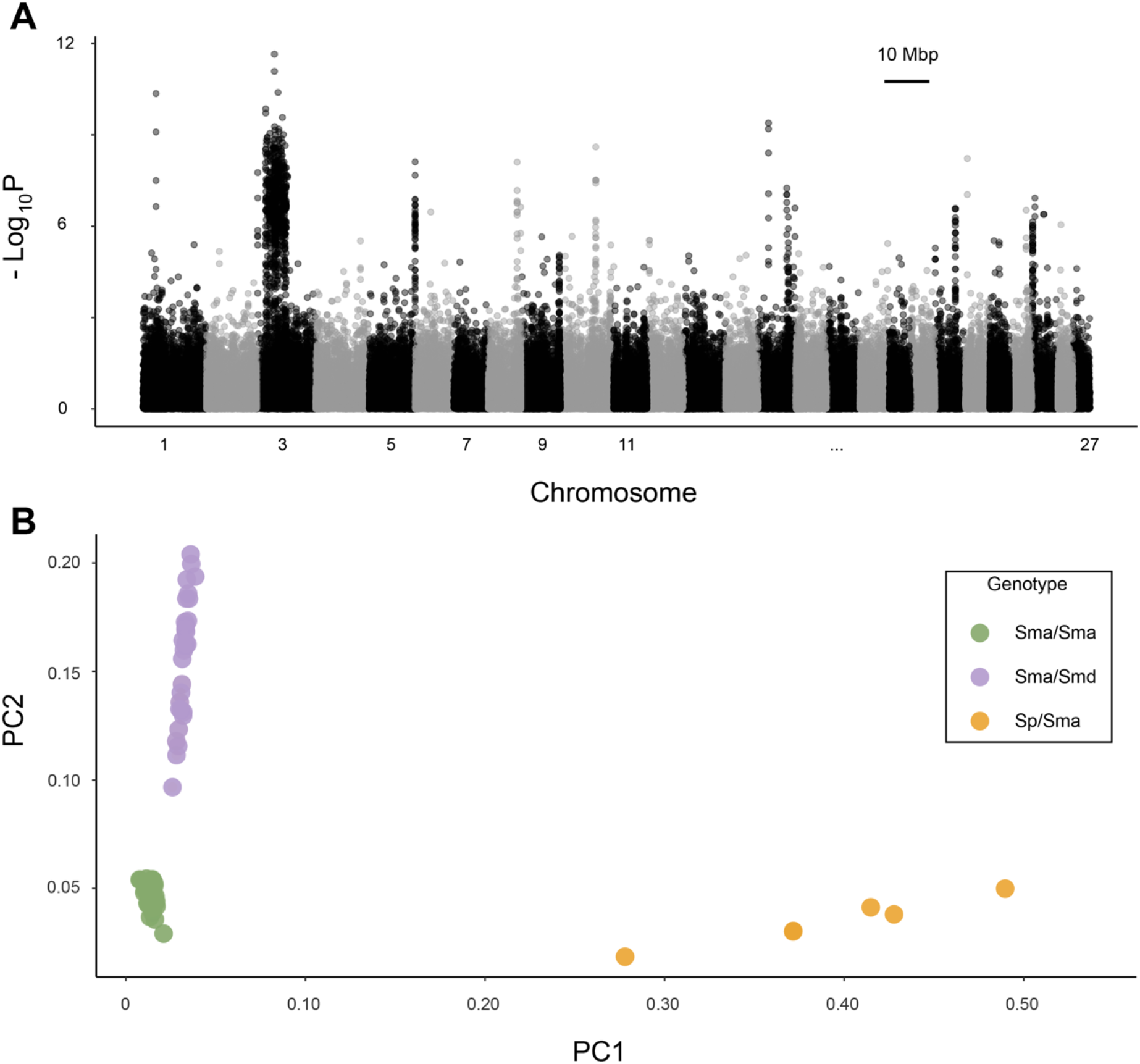
(A) A genome-wide association study reveals a large region on chromosome 3 significantly associated with colony sex ratio in *F. glacialis.* (B) A principal component analysis (PCA) of variants on chromosome 3 identifies three clusters corresponding to three genotypes, Sma/Sma (green), Sma/Smd (purple), and Sp/Sma (yellow).

**Figure 2.**
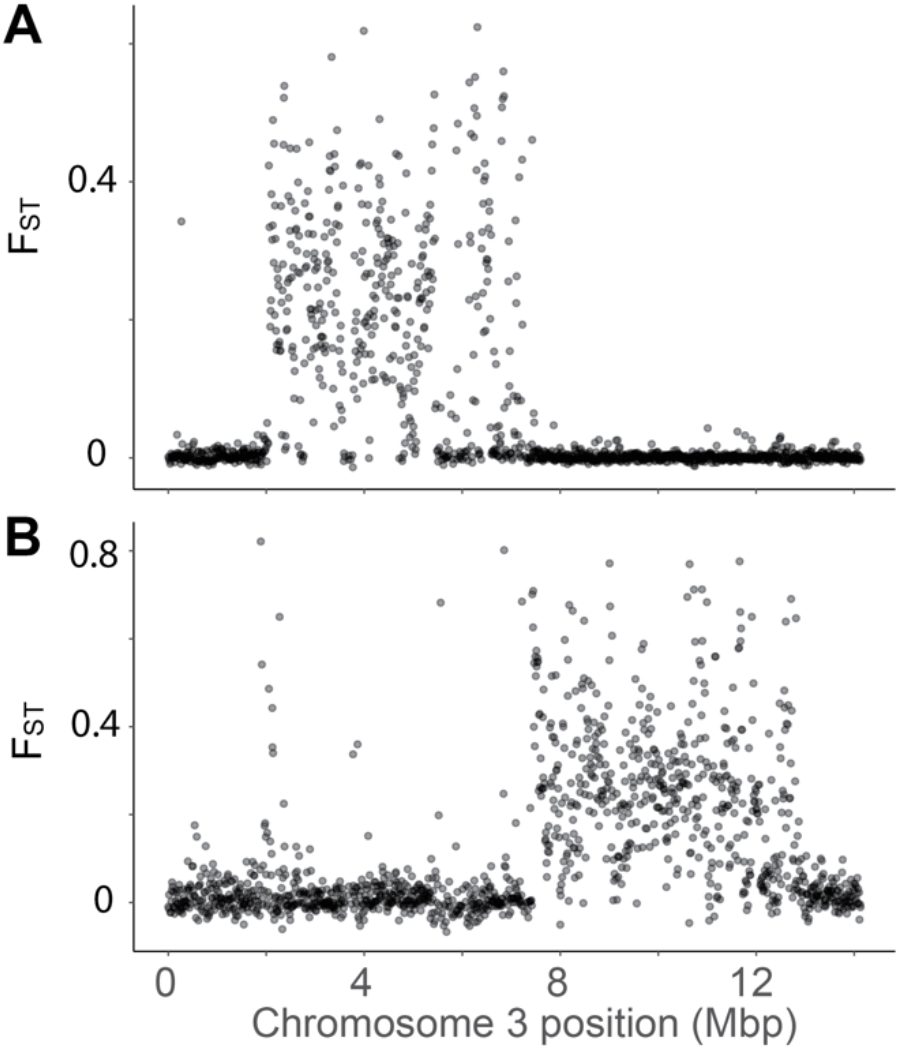
Genetic differentiation between individuals with alternative genotypes on chromosome 3 reveals two adjacent supergenes in *F. glacialis.* (A) Sma/Sma workers and Sma/Smd workers exhibit elevated F_ST_ between 2 Mbp and 7.5 Mbp. (B) Sma/Sma workers and Sp/Sma workers exhibit elevated F_ST_ between 7.5 Mbp and 12.5 Mbp, with a small peak at 2 Mbp. Points represent 10 kbp windows.

Previous studies found that colony queen number in *F. selysi* (28, 57) and other European *Formica* species (29) is controlled by a social supergene on chromosome 3. To determine whether a supergene on chromosome 3 similarly underlies colony queen number in *F. glacialis,* we investigated variation in 19 additional colonies from other populations using ddRADseq. Opposing homozygosity among nestmates (i.e. the presence of two alternative homozygous genotypes in nestmates, which is not possible in haplodiploid full siblings) reveals substantial variation in colony social structure in this species (Fig. 3a), and this variation maps to the supergene region (Fig. 3b). In particular, SNPs that are significantly associated with variation in social structure in the ddRADseq data are localized in the 7.5-12.5 Mbp region (Fig. 3c), corresponding to the region identified in Fig. 2b.

**Figure 3.**
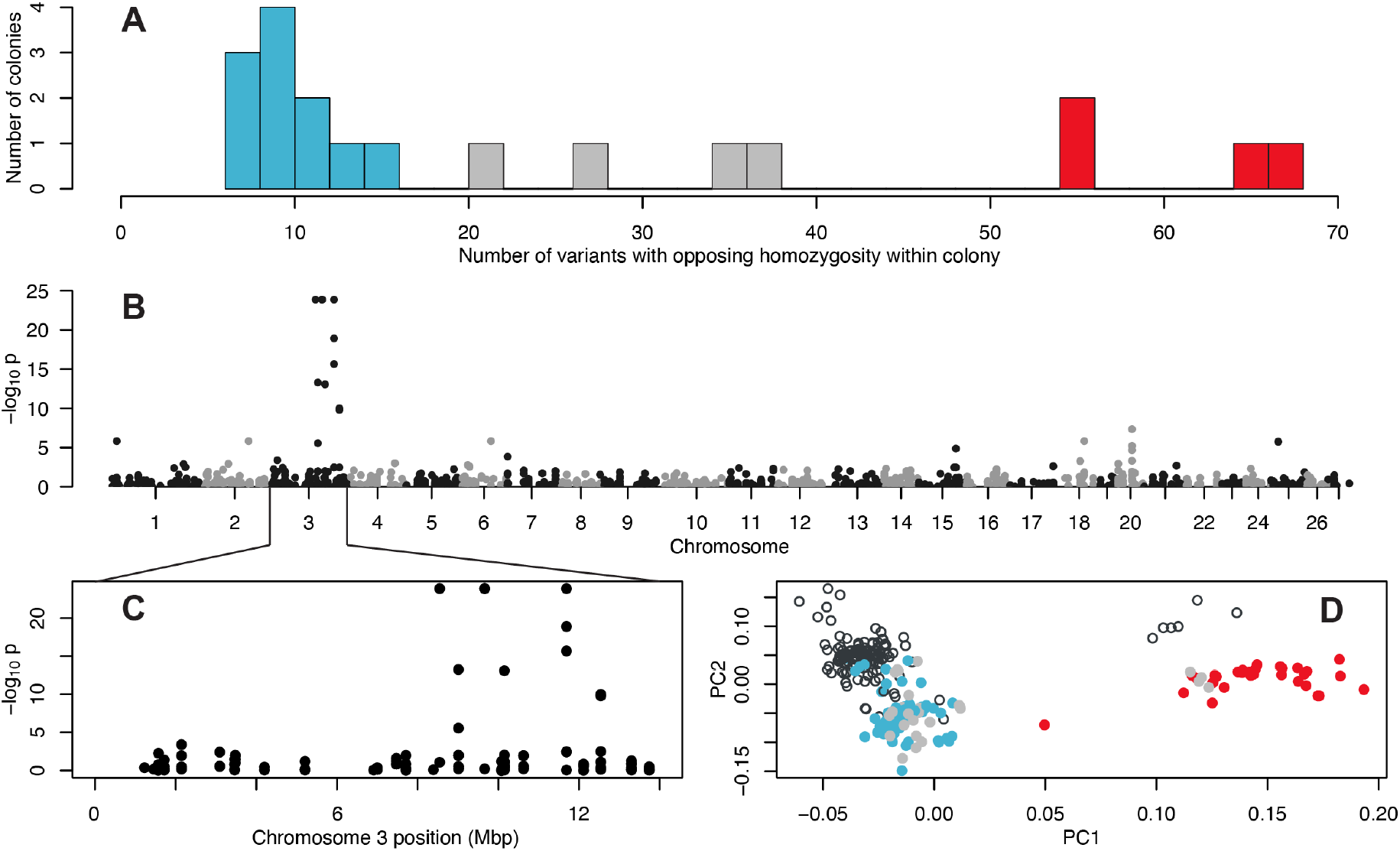
The Sp supergene haplotype is associated with polygyne social structure in *F. glacialis* from Alaska, British Columbia, and Alberta, based on RADseq genotyping of 7-8 workers from each of 19 colonies. (A) Opposing homozygosity varies among colonies. Putative monogyne colonies are colored blue, putative polygyne colonies are colored red, and undetermined are colored gray. (B) GWAS revealed multiple SNPs associated with colony-level opposing homozygosity on chromosome 3. (C) Significantly associated SNPs occur within the 7.5-12.5 Mbp region, also identified in the Yukon population and shown in Fig. 2b. (D) A PCA of variants in both the RADseq (filled circles) and whole genome datasets (open circles) shows that the Sp haplotype identified in the Yukon population clusters with the haplotype associated with polygyne social structure in other populations. The majority of individuals from the Yukon cluster with workers from monogyne colonies in other populations.

A PCA of markers on chromosome 3 that are shared in the whole genome and ddRADseq datasets reveals that the colonies that are assessed to be polygyne based on a high frequency of opposing homozygosity (red, Fig. 3d) consistently harbor one genotype. This genotype is shared with the six individuals from the whole genome sequencing library that formed the yellow cluster in Fig. 1b. These individuals are heterozygous for two alternative supergene haplotypes, one of which appears to occur exclusively in polygyne colonies. We define this haplotype as the Sp haplotype of *F. glacialis.* We note that the Sp found in other *Formica* species spanned about 10.5 Mbp of chromosome 3, from 2 Mbp to about 12.5 Mbp (29), while the one identified here in *F. glacialis* is shorter. The remaining two genotype clusters identified in the whole genome dataset (green and purple clusters, Fig. 1b) both group with workers from colonies assessed to be monogyne in the ddRADseq dataset based on very low levels of opposing homozygosity (Fig. 3a). Based on the regions of differentiation among genotype clusters (Fig. 2), we hypothesized that individuals from the purple cluster carry two alternative supergene haplotypes in the 2-7.5 Mbp region of chromosome 3 (subsequently confirmed with PCR-RFLP genotyping; Fig. 4). One of these haplotypes is found almost exclusively in gyne-producing colonies. The other haplotype is usually homozygous in male-producing colonies. Since one genotype is associated with the production of daughters in monogyne colonies, we name these alleles after mythological twins Danaus and Aegyptus, who had 50 daughters and 50 sons, respectively. Individuals from the gyne-producing cluster (purple, Fig. 1b) have the genotype Sma/Smd, while those from the predominantly male-producing cluster have the genotype Sma/Sma (green, Fig. 1b).

**Figure 4.**
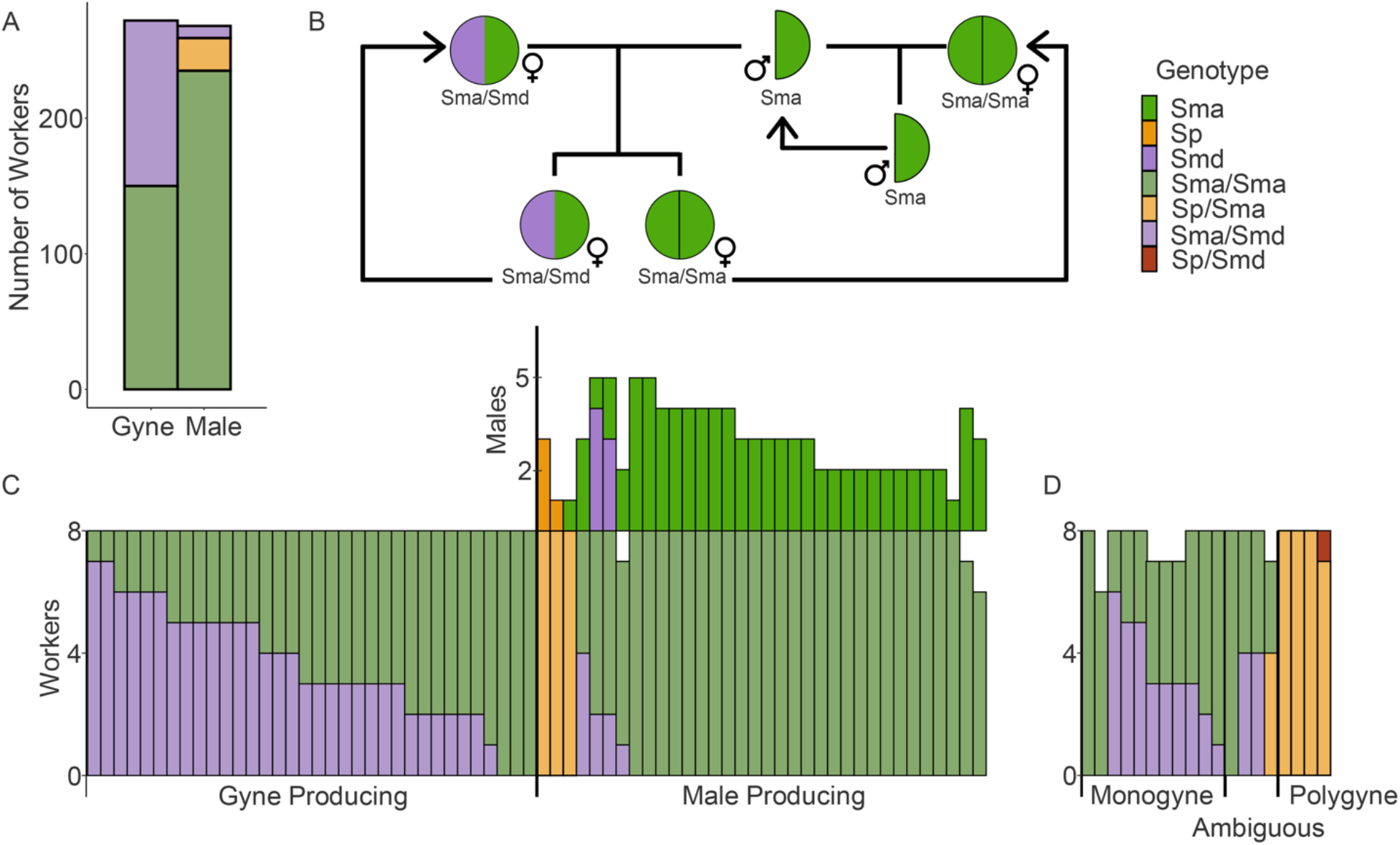
(A) The Sma/Smd genotype in *F. glacialis* occurs in approximately half of workers from gyne-producing colonies, and is rare in workers from male-producing colonies. (B) We propose a model of Mendelian inheritance for maintenance of this supergene system in a largely monogyne population. Heterozygous queens (Sma/Smd) mated with a Sma male produce exclusively female offspring with Sma/Smd and Sma/Sma genotypes. Gynes with the heterozygous genotype become gyne-producers, while homozygotes become male-producers. (C) Gyne-producing colonies usually harbor a mix of Sma/Sma and Sma/Smd workers (31/34 colonies). Male-producing colonies usually contain exclusively Sma/Sma workers and produce Sma males (27/34 colonies). Three additional male-producing colonies contain exclusively Sp/Sma workers and either Sp or Sma males. We did not detect Sma/Smd workers in three gyne-producing colonies, while we found at least one Sma/Smd worker in four male-producing colonies, indicating that the genetic basis of split sex ratio is imperfect in this system. (D) Among monogyne colonies from the broader geographic sample, two contain exclusively Sma/Sma workers, while nine contain Sma/Sma and Sma/Smd workers. All workers from polygyne colonies carry one Sp haplotype. The majority are Sp/Sma, and we detected one Sp/Smd worker.

We developed two PCR-RFLP assays to distinguish these three genotypes in a larger number of individuals from each of the colonies in the focal population in Yukon Territory. Workers from gyne-producing colonies are a mix of Sma/Smd heterozygotes and Sma/Sma homozygotes, while workers from male-producing colonies are most often Sma/Sma homozygotes or Sp/Sma heterozygotes (Fig. 4a). This suggests that gyne-producing monogyne colonies are usually headed by Sma/Smd queens, while male producing monogyne colonies are usually headed by Sma/Sma queens (Fig. 4b). Looking at each colony, we show that 31 out of 34 gyne-producing colonies harbor at least one Sma/Smd worker out of eight genotyped, while 27 out of 34 male-producing colonies harbor only Sma/Sma workers and Sma males (Fig. 4c). Among the remaining male-producing colonies, three harbor only Sp/Sma workers (and are likely polygyne). Of these, two had Sp males and one had a single Sma male. Four male-producing colonies host a mix of Sma/Sma and Sma/Smd workers, as well as both Sma and Smd males. We infer the genotypes of individuals from colonies with known social structure in the ddRADseq dataset using a set of diagnostic SNPs. Across these additional populations, we show that two monogyne colonies harbor exclusively Sma/Sma workers, while nine harbor a mix of Sma/Smd and Sma/Sma workers (Fig. 4d). The four polygyne colonies all contain Sp/Sma workers; one colony contains a single Sp/Smd worker as well.

We obtained a smaller sample of colonies of *F. podzolica,* the sister species of *F. glacialis,* that exhibited a split sex ratio at the focal site in the Yukon Territory. While the GWAS analysis is inconclusive (Fig. S1), we observe similar qualitative patterns in the genomic differentiation between genotype clusters identified in a PCA (Fig. 5). Individuals from these two PCA clusters (Fig. 5a) exhibit elevated genetic differentiation from 2-7.5 Mbp along chromosome 3 (Fig. 5b). Gyne-producing colonies harbor a mix of putative Sma/Smd heterozygotes and Sma/Sma homozygotes. The majority of male-producing colonies contain exclusively Sma/Sma workers (Fig. 5c). A large number of SNPs distinguishing Sma and Smd haplotypes are conserved between *F. podzolica* and *F. glacialis* (Fig. 5d).

**Figure 5.**
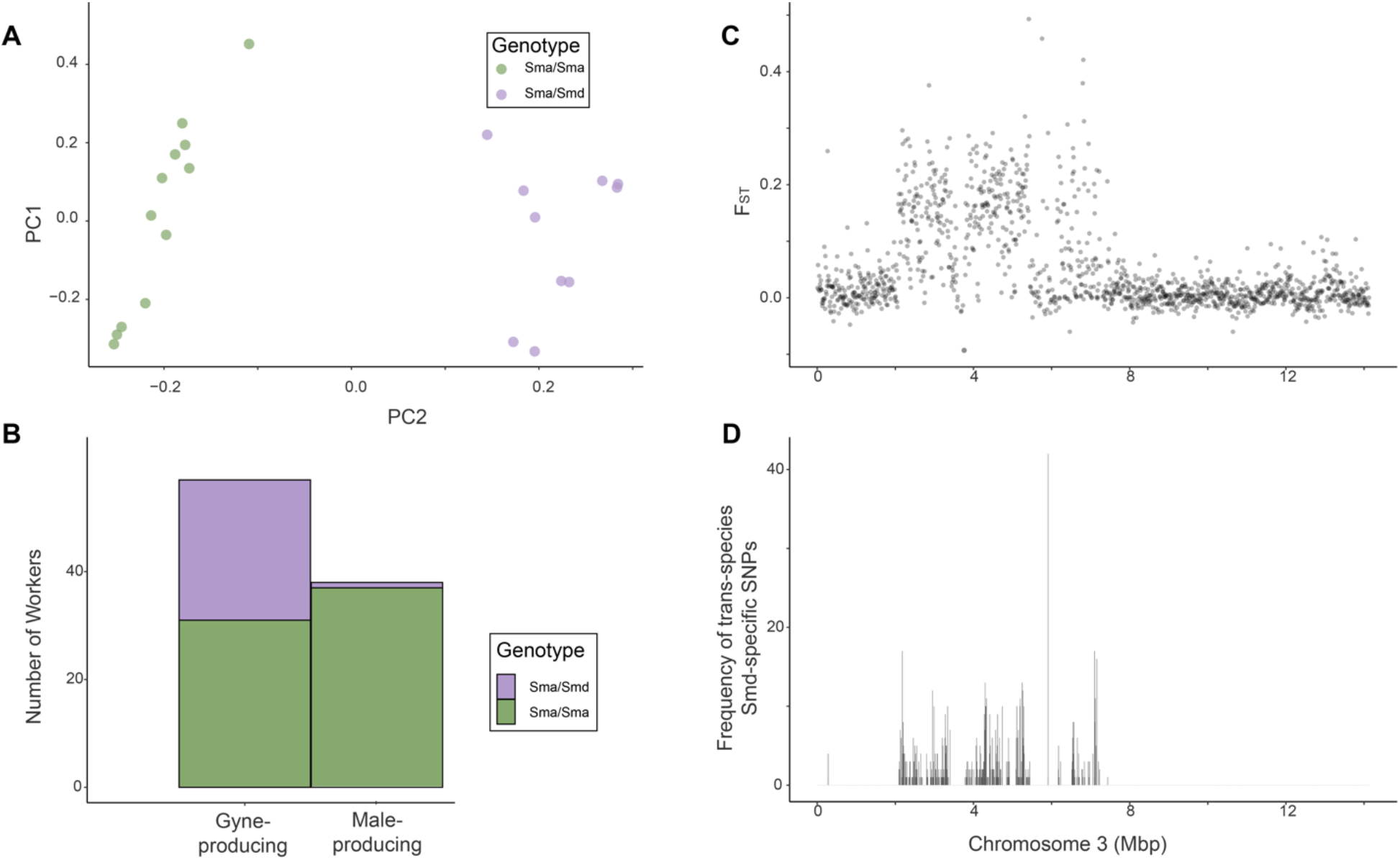
Alternative genotypes matching those detected in *F. glacialis* are also associated with colony sex ratio in *Formica podzolica.* (A) A PCA of genetic markers on chromosome 3 reveals two genotype clusters, Sma/Sma (green) and Sma/Smd (purple). (B) Among workers from gyne-producing colonies, about half carry the Sma/Sma genotype and half carry the Sma/Smd genotype. In contrast, workers from male-producing colonies are almost exclusively Sma/Sma. (C) In *F. podzolica,* the region of elevated F_ST_ between Sma/Sma and Sma/Smd workers spanned 2-7.5 Mbp along chromosome 3. Points represent 10 kbp windows. (D) The *F. glacialis* and *F. podzolica* sex ratio supergenes contain shared SNPs between the putative homologous haplotypes.

## Discussion

We demonstrate that a chromosome underlying queen number across the *Formica* genus is also associated with the split sex ratio patterns observed in a sister species pair. Sex ratio variation based on queen genotype could account for the many empirical exceptions (37) to the patterns predicted by Boomsma and Grafen (36, 38). In *Formica glacialis,* we show that the Smd supergene haplotype behaves like a ‘W’ sex chromosome in that it’s present almost exclusively in females and in a heterozygous state. A key difference is that it influences the sex ratio of offspring rather than the sex of the individual bearer. Single-queen gyne-producing colonies generally harbor a mix of Sma/Smd and Sma/Sma workers, suggesting that the queens are Sma/Smd heterozygotes crossed with Sma males. Through Mendelian inheritance, half of their daughters (the heterozygotes) will in turn be gyne-producing queens, while the other half will be male producers. Males are produced either by homozygous Sma/Sma single queens or by polygyne (Sp/Sma) queens. We noted a few exceptions to this pattern in both gyne- and male-producing colonies, suggesting that genetic control is imperfect. Our focal population was predominantly monogyne, so we do not yet know if some polygyne colonies specialize in producing gynes.

A striking finding of this study is that the extent of the social supergene discovered in other *Formica* species appears to be split into two adjacent, linked supergene regions in *F. glacialis.* One half of the supergene, from 2-7.5 Mbp on chromosome 3, is associated with split sex ratio. The other half, from 7.5-12.5 Mbp, which includes the gene *knockout* identified as a candidate conserved gene influencing social structure in other *Formica* species (29), is associated with social structure (Fig. 2).

These patterns raise questions about the evolution of the functions of these two linked supergenes in *F. glacialis* and *F. podzolica.* Theory predicts split sex ratio to evolve in social hymenopteran populations with variation in relatedness asymmetry. We propose two alternative scenarios that could explain the evolution of these linked regions. In one scenario, we speculate that split sex ratio may have evolved in socially polymorphic *Formica* populations, wherein monogyne and monandrous queens would specialize in gyne production, while polygyne or polyandrous queens would produce predominantly males. Such patterns were documented in other *Formica* species, including *F. truncorum* (42, 58) and Finnish populations of *F. exsecta* (54), although we note that this pattern is not present in all previously studied *Formica* species (46, 51). Specialization in offspring sex ratio based on social structure would select for reduced recombination between loci influencing sex ratio and social structure. In populations with little relatedness asymmetry, as observed in our predominantly monogyne *F. glacialis* population in the Yukon, rare recombinant supergene haplotypes that decouple social determination from sex ratio determination could spread in the population. In this case in particular, we suggest that gene flux from the Sp haplotype onto the Sm may have led to the formation of the Sma haplotype associated with monogyne social structure and the production of males. Such sex ratio supergene systems may persist in species with a mix of socially polymorphic and socially monomorphic populations, which could explain deviations from the theoretical predictions of Boomsma and Grafen (36), wherein predominantly polygyne populations produce highly male biased sex ratios, as in Swiss *F. exsecta* (48), or one social form exhibits strongly split sex ratios and the other is intermediate, as in *F. selysi* (46).

In an alternative scenario, a gene or supergene influencing sex ratio could predate the appearance of persistent social polymorphism; when alternative social structures emerged, selection for male-biased production in colonies with lower average relatedness and for gyne-biased production in colonies with higher average relatedness could have led to the appearance of linked genetic variants favoring one or more queens. The dual roles of linked supergenes in shaping social organization and sex ratio in *Formica* species could help to explain why this supergene has persisted for millions of years (29). Future studies could examine these speculative scenarios by seeking evidence of sex ratio supergenes in other, distantly related *Formica* species. In particular, we predict that a supergene like this one could be particularly evident in stable populations with little relatedness asymmetry.

Our study in *F. glacialis* is not the first to identify sex-specific genetic differences between ant gynes and males (20, 59). However, the mechanisms that produce these sex-specific genetic differences appear to differ across systems, and none are fully understood. Kulmuni and Pamilo (59) showed that hybridization between *Formica aquilonia* and *Formica polyctena* results in admixed females, but that surviving males tend to have a genotype comprised of alleles from only one parental species (59, 60). They proposed that recessive incompatibilities between the genomes of the two species are exposed to selection in haploid males. Subsequent research on this system has revealed instability in the direction of selection on introgressed alleles in males across a ten-year time interval, with introgression recently favored on average in loci where introgression was previously selected against (61). In the tawny crazy ant *Nylanderia fulva,* males invariably carry the same allele at two out of 12 microsatellite loci, while females are almost always heterozygous at these loci (20). Diploid eggs that were homozygous for the male-associated alleles at these loci failed to develop. In *F. glacialis,* we similarly observe a haplotype, Smd, that is found almost exclusively in females. However, we speculate that the mechanism underlying the sex-specific genetic differences in *F. glacialis* may not rely on lethal effects of alleles in one sex or the other, as it appears to in the aforementioned systems. We suggest that the Sma/Smd queens lay exclusively fertilized eggs, which would preclude the production of males. Further research is needed in all systems to compare the specific mechanisms maintaining these genetic differences between the sexes.

Some previous empirical discoveries still need further examination in the light of the newly discovered split sex ratio supergene. Several experimental studies provided evidence that environmental quality and diet can influence colony sex ratio, including in a population of *F. podzolica* from central Alberta (44). We posit that there may be as yet undetected gene x environment interactions, which could explain the rare deviations from expected genotype distributions in workers from our gyne- and male-producing colonies. Repeating food supplementation experiments in colonies of known genetic structure will help to resolve this question. Although the linkage between social and sex ratio supergenes hints at a role of parent-offspring conflict in shaping split sex ratio in *Formica* ancestors (34, 38), many questions remain about how worker control could function in a system with genetic determination of sex ratio. Understanding how the sex ratio supergene functions will help to illuminate how the contemporary conflict plays out. For example, does the Smd haplotype cause cessation of haploid egg production? What factors prevent female offspring of Sma/Sma queens from developing into gynes instead of workers?

Despite limited sampling, we also document an intriguing deviation in the mode of action of the *F. glacialis* social supergene compared to that of *F. selysi.* Across the individuals in the RADseq dataset collected from polygyne colonies, we did not detect any Sp/Sp homozygous individuals (Fig. 4). Polygyne colonies harbored almost exclusively Sp/Sma workers (N = 30), with only a single Sp/Smd worker. In contrast, polygyne *F. selysi* colonies contain exclusively Sp/Sm and Sp/Sp workers and Sp males (28, 33, 57). We did not detect systematic variation at the supergene in monogyne *F. selysi* workers (28, 57), but we note that analyses were carried out with relatively sparse RADseq markers, so it is possible that a small sex ratio supergene could be present in *F. selysi*.

Here, we describe a supergene that drives offspring sex ratio in ants. This sex ratio supergene is closely linked with a previously described supergene that underlies colony queen number in *Formica* ants. The discovery that split sex ratio has a genetic basis helps to resolve the conflicting empirical results about whether and how split sex ratio emerges to resolve parent-offspring conflict in social hymenopterans. We suggest that genetic control of sex ratio should be investigated in other social insects, particularly in those that do not conform to theoretical predictions.

## Materials and Methods

### Sampling and Field Observations

We sampled a mixed population of *F. glacialis* and *F. podzolica* 50 km west of Whitehorse, Yukon Territory, Canada in July 2016. We removed the top 5-10 cm of soil from each nest mound and assessed the presence and sex of winged sexuals. When we observed strongly biased sex ratios (i.e., of the first 10 sexuals examined, at least nine were of the same sex), we sampled at least eight workers and up to five males. We did not sample gynes. In total, we sampled 71 *F. glacialis* colonies, of which 34 were male-producing and 34 gyne-producing. The remaining three sampled colonies contained no *F. glacialis* sexuals; two of the three also contained workers of the socially parasitic species *Formica aserva.* We estimate that at least 80% of colonies with winged sexuals exhibited a biased sex ratio.

### Whole-genome sequencing

We sequenced 138 genomes of workers from 71 *F. glacialis* colonies. We extracted Genomic DNA using the Qiagen DNeasy insect tissue protocol and prepared whole-genome DNA libraries using a low-volume Illumina Nextera protocol (62) with the following modifications: 2 ng/*μ*L input DNA concentration instead of 0.5 ng/*μ*L, tagmentation reaction in 5 *μL* volume instead of 2.5 *μ*L, PCR using Q5 DNA polymerase (New England Biolabs) instead of KAPA HiFi, 90 seconds extension time in the thermal cycling program instead of 30 seconds, and a 0.6:1 ratio of magnetic beads to sample in the magnetic bead clean-up, instead of 1:1 ratio. The libraries were sequenced on an Illumina HiSeq X-Ten by Novogene, Inc., using 150 bp paired-end reads.

### Variant calling

We merged overlapping paired-end reads with PEAR v0.9.10 (63), aligned the reads to the *F. selysi* reference genome (29) using BWA-MEM v0.7.17 (64), and removed PCR duplicates with Samtools v1.8 (65). We called variants using Samtools mpileup v1.8 (66) and filtered the genotypes for missing data (20% per locus, --max-missing 0.8), minor allele count (--mac 2), and minimum depth (--minDP 1) with VCFtools v0.1.13 (67).

### Population genetic analyses

We identified regions significantly associated with colony sex ratio in 71 *F. glacialis* colonies (n=138 individuals) by performing a Genome-Wide Association Study (GWAS) using a univariate linear mixed model implemented in GEMMA v0.94 (68), using a genetic similarity matrix to control for population structure as a random effect. Three colonies without a sex ratio phenotype were assigned an “NA” phenotype. Upon detecting a large region of chromosome 3 significantly associated with sex ratio, we performed a principal component analysis (PCA) using Plink v1.90b3.38 (69) on variants on this chromosome. We calculated Weir and Cockerham’s F_ST_ (70) in 10 kbp windows between the three genotype clusters identified in the PCA using VCFtools v0.1.13 (67).

### *Comparisons with sister species* F. podzolica

We examined the underlying genetics of split sex ratio in the sister species, *F. podzolica*, as well. We sampled 12 colonies (5 male-producing, 7 gyne-producing) from the same Yukon locality, and sequenced the genomes of 22 workers and called variants using the same methods as for *F. glacialis*. We identified genetic clusters based on variants on chromosome 3 and calculated F_ST_ between genetic clusters, again using the methods described above for *F. glacialis.* We identified SNPs with alleles specific to the Smd haplotypes of both species by comparing allele frequencies in four groups: *F. glacialis* Sma/Sma, *F. glacialis* Sma/Smd, *F. podzolica* Sma/Sma, and *F. podzolica* Sma/Smd. Loci with putative Smd-specific alleles shared between both species were defined as those which have allele frequency between 0.4 and 0.6 in both of the Sma/Smd groups, and allele frequency >0.95 or <0.05 in both of the Sma/Sma groups. We plotted the frequency of SNPs meeting these criteria in 10 kbp windows along chromosome 3.

### Social Organization

We sampled 8 workers from 19 additional *F. glacialis* colonies in Alaska, British Columbia, and Alberta, where no winged sexuals were visible at the time of collection. We genotyped 145 of these workers using the double-digest RAD sequencing protocol of Brelsford et al. 2016 (71), with restriction enzymes SbfI and MseI. RAD libraries were sequenced on the Illumina HiSeq 4000 platform by the QB3 Genomics core facility of University of California Berkeley, with 100bp paired-end reads. We aligned reads and called variants using the procedures described above for whole-genome data, but omitting the removal of PCR duplicates. Raw variants were filtered using VCFtools v0.1.13 (67), retaining genotypes with sequence depth of at least 7 and variants with genotype calls in at least 80% of samples.

To assess social organization of the 19 colonies, we calculated the number of loci exhibiting opposing homozygosity within each colony, i.e., at least one worker homozygous for the reference allele and one worker homozygous for the alternate allele. In haplodiploid organisms, a male transmits the same allele to all of his offspring, so in a group of full siblings, opposing heterozygosity is expected to be absent except in the cases of genotyping errors or de novo mutations. Colonies with multiple queens, or with a multiply mated queen, are expected to have a higher number of loci with opposing heterozygosity.

We conducted a genome-wide association study for variants associated with colony-level opposing homozygosity using a linear mixed model implemented in Gemma v0.94 (68), which uses a relatedness matrix to control for non-independence of samples. Finally, we carried out a principal component analysis of variants on chromosome 3 on a merged dataset of whole-genome and ddRAD *F. glacialis* genotypes. We generated a list of variants on chromosome 3 present in both the whole-genome and ddRAD filtered VCF files, extracted those variants from both datasets, and generated a merged VCF using VCFtools v0.1.13 (67). We used Plink v1.90b3.38 (69) to carry out a principal component analysis on the resulting merged genotypes.

### Colony genotype distributions

We designed a targeted PCR-RFLP assay for a trans-species SNP tagging alternative chromosome 3 haplotypes in both *F. glacialis* and *F. podzolica.* We designed primers (CTGGAACAACGGATCCTCA and TTCGCGATTCGAATTTCTC) to amplify a 338 bp fragment, which, when digested with the restriction enzyme MluCI, produces fragments of 325 and 13 bp for the haplotype associated with gyne production and 223, 102, and 13 bp for the haplotype associated with male production. We used this assay to genotype 6 additional workers and any available males for all colonies of both species.

We designed a second PCR-RFLP assay for a trans-species SNP broadly conserved across Formica that differs between Sm and Sp alleles of the gene *knockout.* Primers GGTGGYTCTTTCAACGACG and GCCATGTTCACCTCCACCA amplify a 230 bp fragment, which when digested with the restriction enzyme HinfI produces fragments of 132 and 98 bp for the Sm allele and 230 for the Sp allele. We used this assay to genotype 6 additional workers and any available males from three *F. glacialis* colonies where initial whole-genome sequencing of two workers identified the presence of the Sp allele at *knockout.* For both PCR-RFLP assays, we visualized the distinct banding patterns with 2% agarose gel electrophoresis.

We constructed bar plots of supergene genotype frequency by colony based on whole-genome sequencing and PCR-RFLP genotyping for the Yukon population, and based on ddRAD for Alaska, Alberta, and British Columbia populations.

## Data Sharing

Raw sequences will be uploaded to NCBI SRA (accession numbers to be determined pending acceptance). Colony metadata including locality, observed sex ratio, and inferred social structure is included in Dataset S1.

## Acknowledgements and funding sources

This material is based upon work supported by the National Science Foundation Graduate Research Fellowship to GLR under Grant No. DGE-1326120, by NSF DEB grant 1733437 to AB and JP, and by U. S. Department of Agriculture National Institute of Food and Agriculture Hatch #CA-R-ENT-5126-H to JP. This work used the Vincent J. Coates Genomics Sequencing Laboratory at UC Berkeley, supported by NIH S10 OD018174 Instrumentation Grant. The authors thank K. Martinez, Z. Alam, and A. Klement for their assistance in the lab.

---

**Fig. S1.**
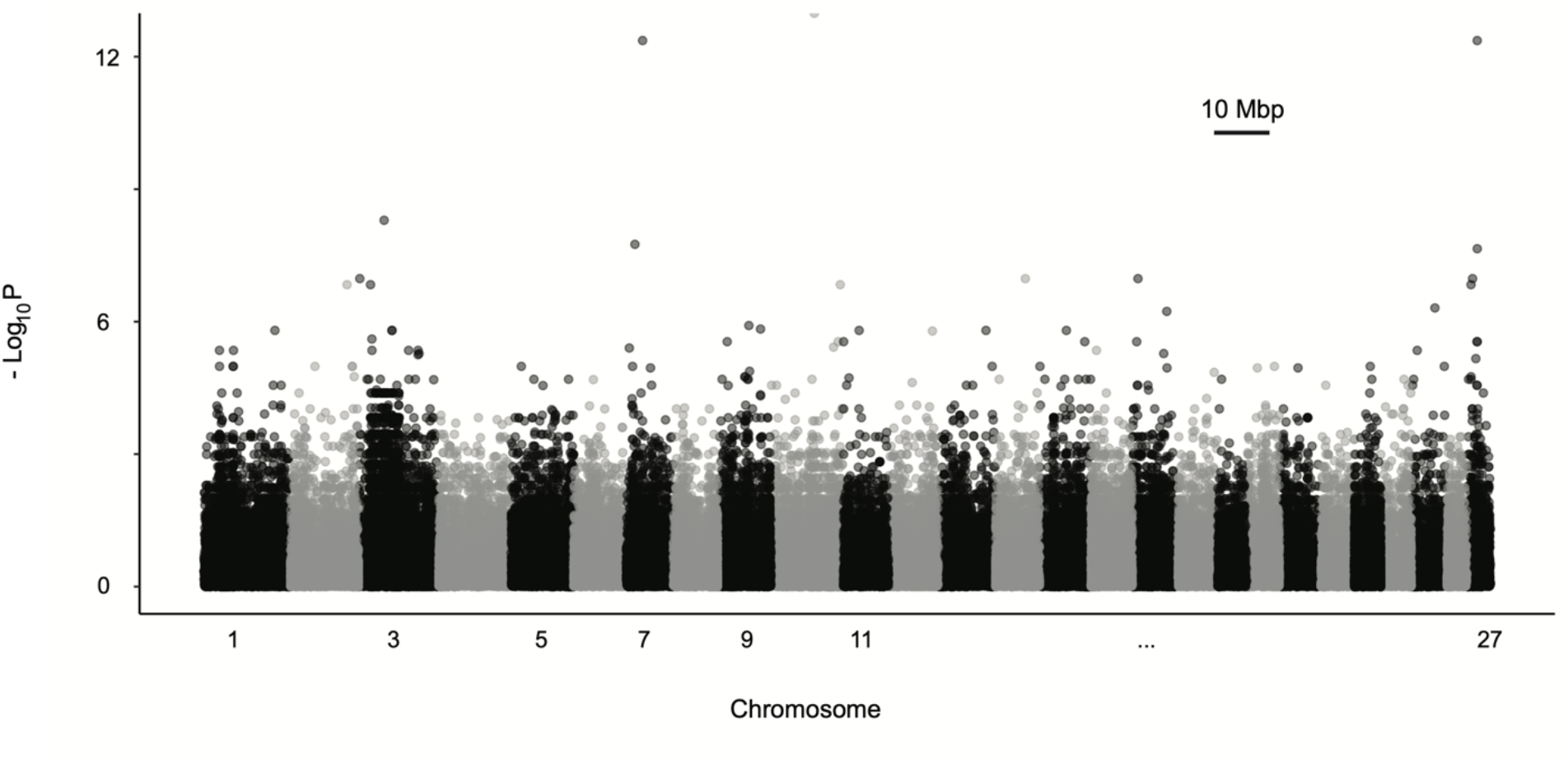
Manhattan plot of GWAS results in *Formica podzolica* shows no clear association between colony sex ratio and chromosome 3 supergene region based on 22 sequenced workers from seven gyne-producing and five male-producing colonies.

